# Development and Characterization of Recombinant Vesicular Stomatitis Virus (rVSV)-based Bivalent Vaccine Against COVID-19 Delta Variant and Influenza Virus

**DOI:** 10.1101/2021.12.14.472657

**Authors:** Zhujun Ao, Maggie Jing Ouyang, Titus Abiola Olukitibi, Bryce Warner, Robert Vendramelli, Thang Truong, Manli Zhang, Sam Kung, Keith R Fowke, Darwyn Kobasa, Xiaojian Yao

## Abstract

COVID-19 and influenza are both highly contagious respiratory diseases with a wide range of severe symptoms and cause great disease burdens globally. It has become very urgent and important to develop a bivalent vaccine that is able to target these two infectious diseases simultaneously. In this study, we generated three attenuated replicating recombinant VSV (rVSV) vaccine candidates. These rVSV-based vaccines co-express SARS-CoV-2 Delta variant spike protein (SP) or the receptor binding domain (RBD) and four copies of the highly conserved M2 ectodomain (M2e) of influenza A fused with the Ebola glycoprotein DC-targeting/activation domain. Animal studies have shown that immunization with these bivalent rVSV vaccines induced efficient but variable levels of humoral and cell-mediated immune responses against both SARS-CoV-2 and influenza M2e protein. Significantly, our vaccine candidates induced production of high levels of neutralizing antibodies that protected cells against SARS-CoV-2 Delta and other SP-pseudovirus infections in culture. Furthermore, vaccination with the bivalent VSV vaccine via either intramuscular or intranasal route efficiently protected mice from the lethal challenge of H1N1 and H3N2 influenza viruses and significantly reduced viral load in the lungs. These studies provide convincing evidence for the high efficacy of this bivalent vaccine to prevent influenza replication and initiate robust immune responses against SARS-CoV-2 Delta variants. Further investigation of its efficacy to protect against SARS-CoV-2 Delta variants will provide substantial evidence for new avenues to control two contagious respiratory infections, COVID-19 and influenza.

## Introduction

The ongoing pandemic of coronavirus disease 2019 (COVID-19) has been the most serious threat to global public health, with a total number of cases surpassing 256 million and over 5 million deaths ^1, 2^. The case fatality ratio (CFR) of COVID-19 is 2.0% as of 18 November 2021, which is close to the CFR of 1918 flu (>2.5%) and shockingly higher than other influenza pandemics (<0.1%) ^3-5^. Since the identification of SARS-CoV-2 sequences ^6^, extensive worldwide efforts have focused on developing effective vaccines and antiviral drugs against SARS-CoV-2. Several vaccines have been successfully developed and approved for the prevention of COVID-19 ^7^. However, the spread of some highly transmissible variants of concern (VOCs) and their ability to infect immunized people (breakthrough infections) have challenged the effectiveness of current vaccines. This raised a debate about the need for reformulated vaccines targeting these VOCs.

SARS-CoV-2 is a member of the betacoronavirus subfamily that causes severe symptoms in respiratory, gastrointestinal and neurological systems ^8-11^. Since late 2019, when the COVID-19 outbreak emerged in Wuhan, China, the COVID-19 pandemic continually threatened public health with the continuing emergence of different VOCs. The mutation D614G (amino acid (aa) 614 from aspartic acid to glycine) in the SP receptor-binding domain (RBD) was the first mutation found to endow the virus with a higher transmissibility, indicating that more virulent strains may emerge due to the fast evolution of the virus ^12, 13^. One of the most virulent and highly transmissible VOC strains is the Delta variant (B.1.617.2). The Delta variant of SARS-CoV-2 was first found in India in Dec. 2020, and only in several months did this particular variant spread worldwide, becoming the dominant variant in many countries, including India, the U.K., Israel and the United States ^14^. To date, the Delta variant has been the most contagious of all the known SARS-CoV-2 variants. A recent study found that people infected by the Delta variant had viral loads that increased to more than 1000 times higher than those of individuals infected with the original strain in 2020 ^15^. Therefore, it is imperative to develop a vaccine that can specifically target the Delta variant to more efficiently block the transmission and infection of SARS-CoV-2 worldwide.

Influenza is another contagious respiratory illness caused by the influenza virus. Surprisingly, 100 years after the pandemic of influenza A virus (IAV) that killed approximately 50 million people globally in 1918 ^16, 17^, seasonal influenza still poses a large threat to public health, with a global annual mortality of over 300,000 ^18^. Vaccination remains the most effective method to prevent influenza-associated illness; however, the effectiveness of the seasonal influenza vaccine is only approximately 10% to 60% since the vaccine strains may not be well matched to circulating strains ^19, 20^. Therefore, it is necessary to develop a universal influenza vaccine that can elicit immune responses against all influenza strains regardless of the virus subtype, antigenic drift or antigen shift. Recently, we demonstrated that recombinant rVSV-EΔM-tM2e which contains four copies of the highly conserved extracellular domain of the influenza matrix protein (M2e) could efficiently protect mice from influenza H1N1 and H3N2 challenges ^21^. Given that both COVID-19 and influenza are contagious respiratory diseases mainly transmitted during the same seasons with an increasing threat to the globe, it is necessary to develop a multivalent vaccine that could simultaneously protect against both COVID-19 and influenza.

Vesicular stomatitis virus (VSV) is a single-stranded negative-sense RNA virus belonging to the family *Rhabdoviridae*. Although VSV can cause illness in livestock and some other animals, it is highly restricted in humans by the IFN response and generally causes no or little symptoms in humans ^22^. The VSV-based vaccine platform has been used as an attenuated replication-competent vaccine that induces a rapid and robust immune response to viral antigens after a single immunization and has been shown to protect against several pathogens, including Ebola virus, Zika virus, HIV, and Nipah virus ^23-27^. Specifically, the VSV-based Zaire Ebola glycoprotein (GP) vaccine (rVSV-ZEBOV) that expresses EBOV GP induced robust and persistent specific antibodies against EBOV ^27, 28^ and was considered safe and effective against EBOV in a phase III clinical trial. Intriguingly, a recent report indicated that intranasal vaccination with VSV-SARS-CoV-2 resulted in protection in hamsters if administered within 10 days prior to SARS-CoV-2 challenge, and that animals did not show signs of pneumonia, demonstrating that VSV-based vaccines are fast-acting vaccine candidates that are protective against COVID-19.

In this study, we generated several rVSV bivalent vaccine candidates that co-expressed SARS-CoV2 Delta variant spike protein (SP) or RBD and four copies of highly conserved influenza M2 ectodomain (M2e) fused with a DC-targeting/activation domain derived from EBOV GP (EboGPΔM) based on our previously reported vaccine platform^21, 29^. Here, we characterized the expression of SARS-CoV-2 Delta variant spike protein (SP) or RBD and influenza M2 ectodomains of these bivalent vaccine candidates and their abilities to induce immune responses against SARS-CoV-2 SP, especially Delta SP, and influenza M2e. The study demonstrated that vaccination with the bivalent VSV vaccine efficiently protected mice from lethal challenges of influenza H1N1 viruses.

## Results

### Generation of rVSV-based vaccines expressing both the conserved M2 ectodomain (M2e) of influenza and SARS-CoV-2 Delta spike protein

Given that a new SARS-CoV-2 Delta variant (B1.617.2) has become a dominant variant spreading rapidly around the world ^14^, it is necessary to develop a vaccine that specifically targets the Delta strain and related variants. To this end, we generated several bivalent rVSV-based vaccines against both SARS-CoV-2 Delta variants and influenza virus. First, we performed cDNA synthesis and two step PCR to generate the cDNAs encoding SARS-CoV-2 Delta variant spike protein (SP_Delta_) containing a C-terminal 17 amino acid (aa) deletion (SPΔC) (Fig. 1A, a, Suppl. Fig. 1). The deletion of 17 aa at the C-terminus of SP will facilitate the transportation of SP to the plasma membrane and its assembly into virus because the assembly of SARS-CoV-2 occurs in the ER-Golgi intermediate compartment ^30, 31^. To reduce the cytotoxic effect of SPΔC_Delta_ ^32^ in the vaccine platform, we also introduced an isoleucine (I) to alanine (A) substitution at position 742 aa and named it SPΔC1 (Fig. 1A, a). The analyses revealed that the I742A point mutation in SPΔC1 significantly reduced pseudovirus infectivity (Fig. 1B) and syncytia formation compared to SPΔC_Delta_ in A549-ACE2 cells (Fig. 1C and D). In SP_Delta_ΔS2ΔC (named SPΔC2), a 381 aa fragment in the S2 domain (744-1124 aa) was further deleted (Fig. 1A, b). Meanwhile, we inserted a receptor-binding domain (RBD) from SARS-CoV-2 (wild type) SP into the Ebola glycoprotein (EboGPΔM) to replace the mucin-like domain (MLD) and named it EboGPΔM-RBD (ERBD) (Fig. 1A, c). Finally, we inserted cDNA encoding SPΔC1, SPΔC2 and ERBD into a recently reported rVSV-EM2e vaccine vector, which contains an EboGPΔM fused with four copies of influenza M2 ectodomain (24 aa) polypeptide (EboGPΔM-tM2e, or EM2) ^21^ (Fig. 1A, d) and named them V-EM2e/SPΔC1, V-EM2e/SPΔC2, and V-EM2e/ERBD (Fig. 1D).

**Figure 1.**
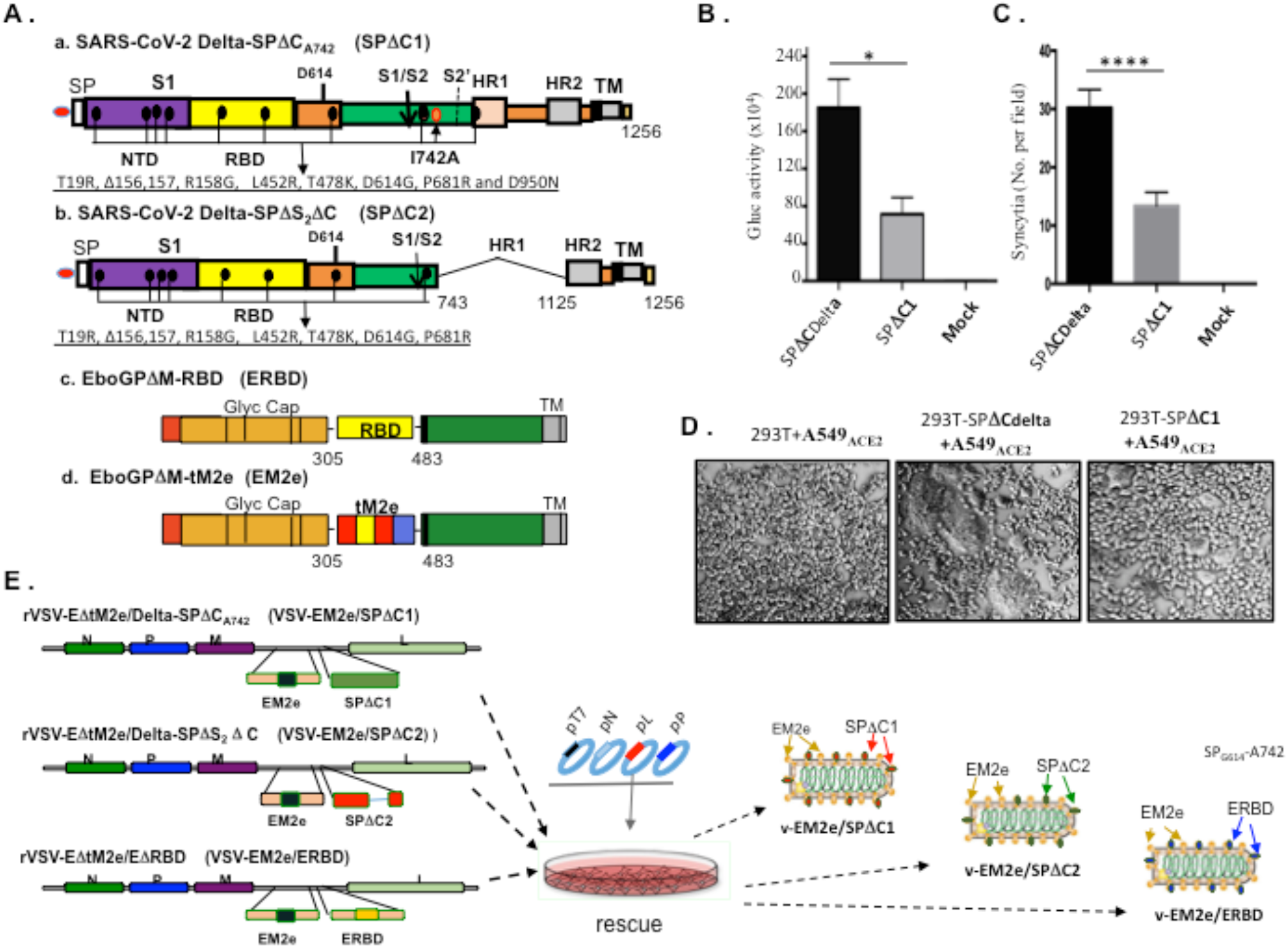
Construction and rescue of rVSV Delta SP and influenza M2e bivalent vaccines. A) Schematic diagram of the Delta SPΔC and EboGPΔM-tM2e immunogens present in the bivalent vaccines. a. SARS-CoV-2 Delta-SPΔC_A742_ (SPΔC1), containing a C-terminal 17 aa (DEDDSEPVLKGV<u>KLHYT</u>) deletion and a I742A mutation as indicated. The nine mutations in Delta SP are listed in lower part. b. Delta SPΔC2, containing the C-terminal 17 aa deletion and another 381 aa (encompassing aa744 to aa1124) deletion in S2 domain. The eight mutations in SPΔC2, are listed in lower part. c. EboGPΔM-RBD, the RBD of SARS-CoV-2 was used to replace the MLD domain in EboGP. d. EboGPΔM-tM2e, four copies of influenza virus M2 ectodomain (24 aa) polypeptide (tM2e) replaced the MLD domain in EboGP. B) The attenuated virus entry of SPΔC1. A549_ACE2_ cells were infected with equal amounts of SPΔC_Delta_-PVs or SPΔC1-PVs (adjusted by P24) carrying Gaussia luciferase (Gluc) gene, as indicated. At 48hrs after infection, the Gluc activity in the supernatant of different infected cultures was measured. Data represents Mean ±SD of two replicates from a representative experiment out of three performed. C and D) The attenuated cell-to-cell fusion ability of SPΔC_Delta_- or SPΔC1-mediated syncytia formation was analyzed by co-culturing the SPΔC_Delta_- or SPΔC1-expressing 293T cells with A549_ACE2_ cells. The amounts of syncytia were counted after 24 hrs in 5 different views of microscope (C), and was also imaged under bright-field microscopy (D). E) Schematic diagram of VSV-EM2e/SPΔC1, VSV-EM2e/SPΔC2 and VSV-EM2e/ERBD and the virus rescuing procedures. 293T and Vero E6 co-culture cells were co-transfected with VSV-ΔG-EM2/SPΔC1, VSV-ΔG-EM2/SPΔC or VSV-ΔG-EM2/RBD, and helping plasmids (T7, N, L, P plasmids). The supernatants containing V-EM2e/SPΔC1, V-EM2e/SPΔC2 and V-EM2e/ERBD viruses were used to infect Vero E6 cells to generate the rVSV stocks.

The established reverse genetics technology ^33^ was used to rescue three rVSV vaccine candidates. Primary recovery was performed in 293T and VeroE6 co-culture cells by co-transfecting the rVSV-EM2e/SPΔC1, rVSV-EMe2/SPΔC2 or rVSV-EM2e/ERBD vector with the VSV accessory plasmids encoding VSV-N, P, L, and T7 ^34^ (Fig. 1C). After 48 hrs and 72 hrs, the supernatants containing the recovered rVSVs were collected and used to infect Vero E6 cells, which consequently showed a cytopathic effect (CPE) at 72 or 96 hrs post-infection (Fig. 2A). To verify the expression of SPΔC1, SPΔC2, ERBD, and EM2e in each corresponding rVSV-infected Vero E6 cell line, we performed an immunofluorescence assay with a rabbit anti-SARS-CoV-2 RBD antibody or anti-influenza M2 antibody. The results confirmed the expression of SPΔC1, SPΔC2, ERBD and EM2 in the corresponding rVSV-infected cells but not in mock-infected cells (Fig. 2B). Meanwhile, we investigated the expression of the above proteins in the infected cells by western blotting (WB) with the corresponding antibodies. The results showed the abundant expression of full-length SP_Delta_ΔC and cleaved S1 in V-EM2e/SPΔC1-infected cells (Fig. 2C, top panel, Lane 3), while SPΔC2 and ERBD were primarily detected as single bands (Fig. 2C, top panel, Lanes 2 and 4). As expected, the expression of EM2e and VSV nucleocapsid (N) protein was detected in all rVSV-infected cells (Fig. 2C, middle and lower panel, Lanes 2 to 4).

**Figure 2.**
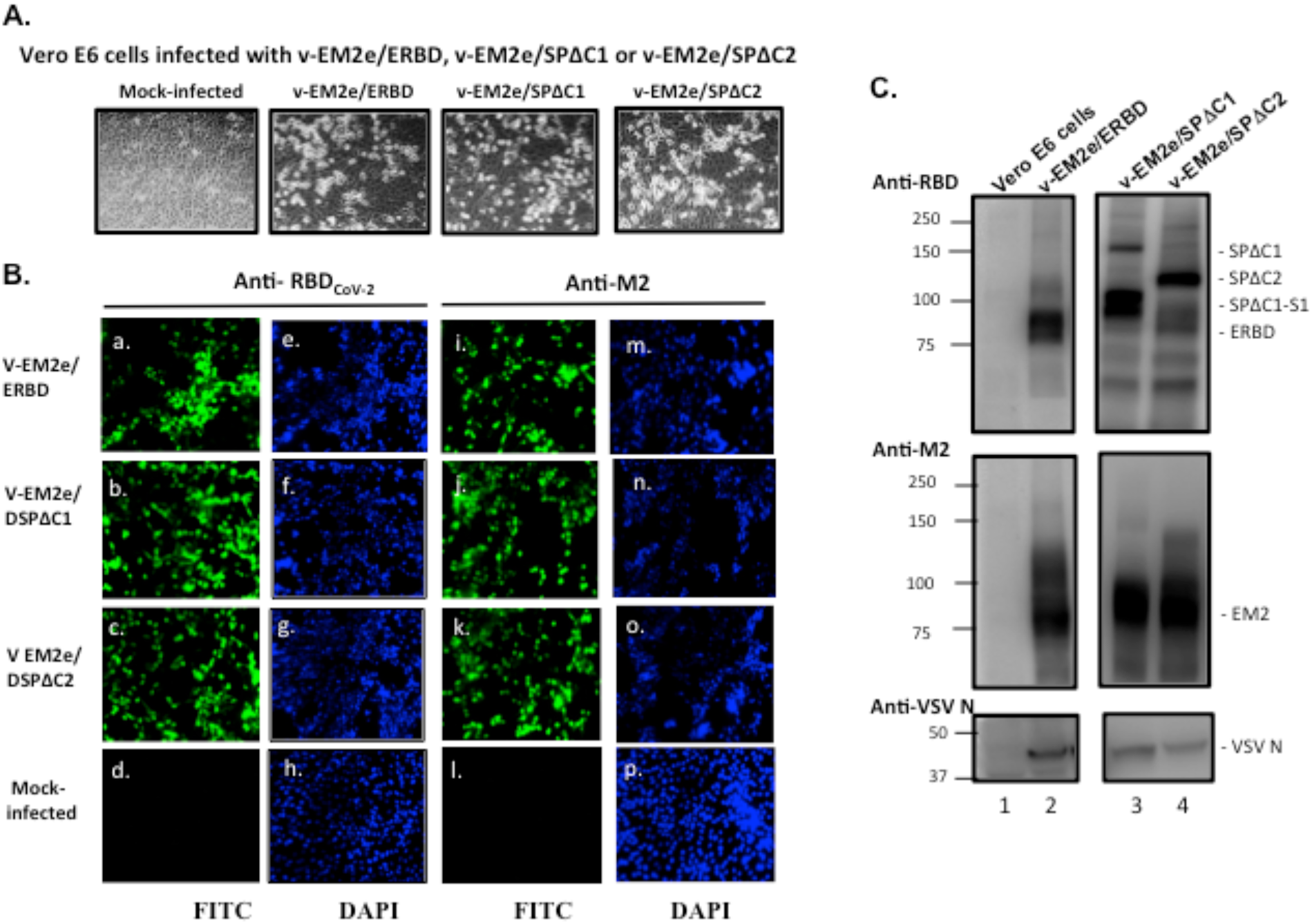
Expression of V-EM2e/SPΔC1, V-EM2e/SPΔC2 or V-EM2e/ERBD in infected VeroE6 cells. A) The infection of V-EM2/SPΔC1, V-EM2/SPΔC2 or V-EM2/ERBD in Vero-E6 cells induced the cytopathic effects after four days of infection. B) Representative immunofluorescence images of Vero E6 cells infected with V-EM2e/SPΔC1, V-EM2e/SPΔC2, V-EM2e/ERBD or mock infected, stained with anti-SARS-CoV-2 RBD antibody (a-d) or anti-M2e antibody (i-l), and DAPI (e-h, m-p). C) VeroE6 cells infected with the rescued V-EM2/SPΔC1, V-EM2/SPΔC or V-EM2/ERBD were lysed and processed with SDS-PAGE followed by WB with a rabbit anti-SARS-CoV-2 NTD antibody (top panel), a mouse antibody against influenza M2e (middle panel) or anti-VSV nucleocapsid (N) (low panel).

### Replication attenuation and different cell tropisms of bivalent VSV vaccine candidates compared to wild-type VSV

In our rVSV vaccine strategy, VSV-G was replaced by EM2e and SPΔC or ERBD, which attenuated the pathogenicity of rVSV. Considering that rVSV is a replication-competent vector, it is necessary to investigate the replication ability and cell tropism of the above rVSV vaccine candidates. We therefore used a dose of 100 TCID50 to infect following cell lines: A549, a type II pulmonary epithelial cell line ^35^; MRC-5, a human fetal lung fibroblast cell line ^36^; U251MG, a glioblastoma cell line; CD4+ Jurkat T cells; human monocyte-derived macrophages (MDMs) and dendritic cells (DCs) (Fig. 3). We assessed the cytopathic effect (CPE) of rVSV vaccine candidates and assessed their growth kinetics. A comparison study revealed that 1) rVSV vaccine candidates were unable to infect CD4+ Jurkat T cells and MRC-5 cells, while wild-type VSV replicated efficiently in all tested cells and induced typical CPEs, such as cell rounding and detachment (Fig. 3A, B). 2) In A549 cells, U251 cells, MDMs and MDDCs, three rVSV vaccine candidates displayed positive infection but exhibited much slower replication kinetics and no significant CPE was observed compared to wild-type VSV during the testing period. This implies that these rVSV vaccine candidates have less replication ability and cytopathic effects. All of these data provided evidence supporting that the replication ability of these replicating rVSV vaccine candidates is highly attenuated compared to wild-type VSV.

**Figure 3.**
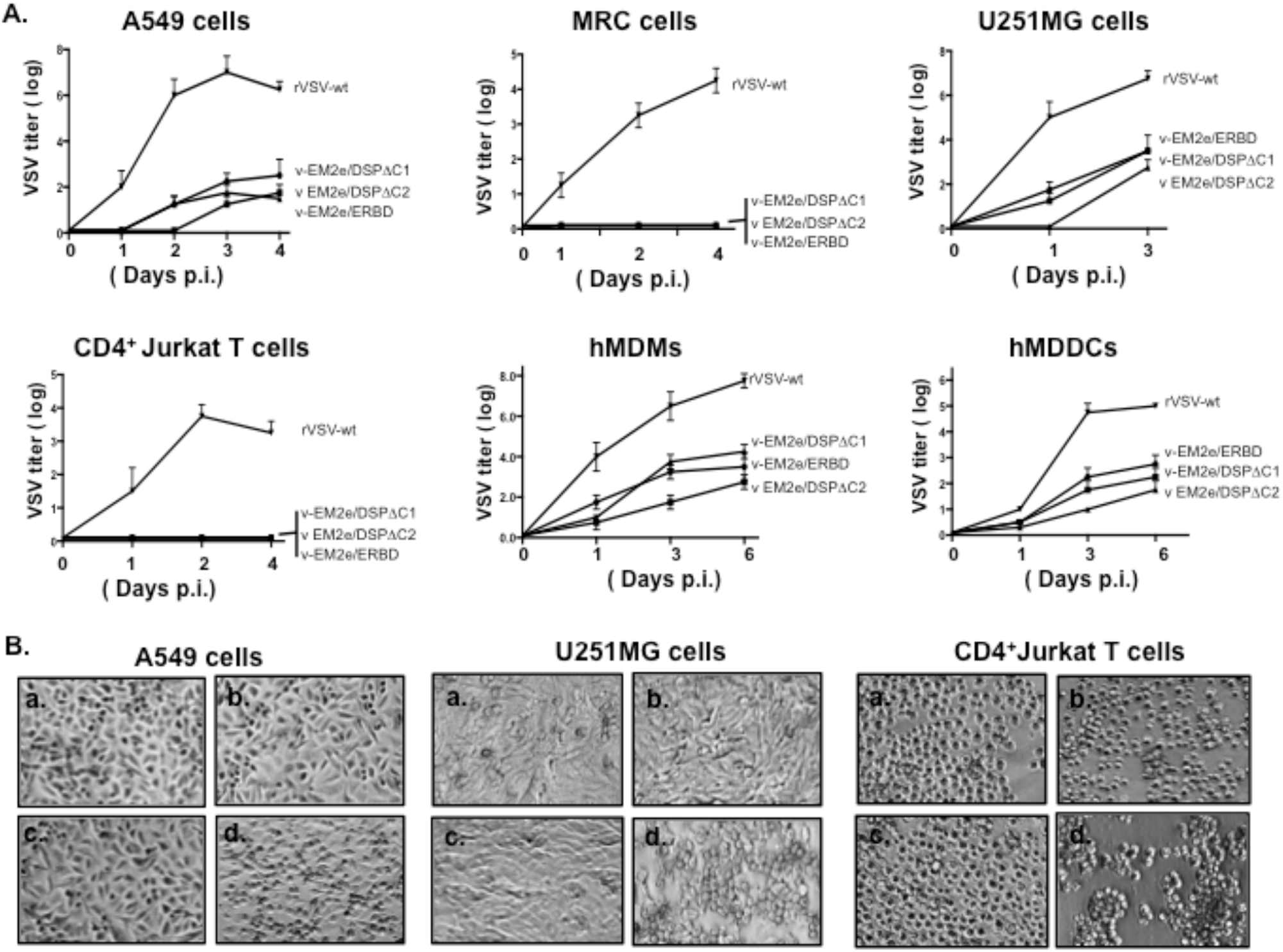
Characterization of the replication kinetics and the cell tropisms of bivalent rVSV vaccine candidates. **A)** Each of bivalent VSV vaccine candidates or the rVSV expressing VSV-G (rVSV-wt) was used to infect different cell lines, including A549, MRC-5, U251MG, CD4^+^ Jurkat T cells, human monocyte-derived macrophages (MDMs) and Dendritic cells (MDDCs). Supernatants were collected at different time points post infection as indicated and were titrated on Vero E6 cells. Data represent Mean ±SD and were obtained from two replicates of a representative experiment out of two performed. B) The ability of induced cytopathic effects in A549, U251MG and CD4^+^ Jurkat T cells, by each rVSV were observed after 4 days of infection under microscopy. a, V-EM2e/ERBD; b, V-EM2e/SPΔC1; c, V-EM2e/SPΔC2, and d. rVSV-wt.

### Evaluation of anti-SARS-CoV-2 and anti-influenza M2 humoral immune responses induced by different bivalent VSV vaccine candidates

To test whether the above bivalent VSV vaccine candidates could induce specific immune responses against SARS-CoV-2 and influenza M2 simultaneously, we administered two doses of immunization (prime on Day 0 and boost on Day 14) in Balb/c mice with each bivalent VSV vaccine candidate intramuscularly (IM, 1×10^8^ TCID_50_) or intranasally (IN, 1×10^5^ TCID_50_). The potential adverse effects and the body weight of the mice were monitored weekly. No changes were noticed clinically, and the steady weight gain of the vaccinated mice was comparable to that of the PBS group (data not shown). Sera from immunized mice were collected on Days 14 and 28 post immunization, and the anti-SARS-CoV-2 RBD and anti-M2 antibody levels were measured using the corresponding antigen-coated ELISA as described in the Materials and Methods. The results showed that 1) IM immunization with V-EM2e/SPΔC1 or V-EM2e/SPΔC2 induced higher levels of anti-SARS-CoV-2 RBD IgG and IgA antibodies than IN immunization (Fig. 4B-E). 2) V-EM2e/ERBD IM administration induced much lower levels of anti-SARS-CoV-2 RBD IgG antibodies than the other two rVSV vaccines (Fig. 4B and C). 3) All vaccine candidates elicited high levels of anti-M2-specific IgG and IgA antibodies regardless of the route of administration (Fig. 4F-I). 4) IN immunization resulted in more complicated anti-RBD and anti-M2 antibody profiles. IN immunization with V-EM2/SPΔC1 induced slightly higher levels of anti-SARS-CoV-2 IgG and IgA antibodies than immunization with V-EM2/SPΔC2 (Fig. 4, B-E, compare bars 3 to 4). Conversely, the levels of anti-M2 IgG and IgA antibodies induced by V-EM2/SPΔC1 were lower than those induced by rVSV-EM2/SPΔC2 (Fig. 4, compare F-I, compare bars 3 to 4). The underlying mechanism(s) is still unclear. Nevertheless, all of these observations indicate that IM- and IN-immunizations with V-EM2/SPΔC1 and V-EM2/SPΔC2 induced efficient anti-SARS-CoV-2 RBD and anti-M2 immune responses.

**Fig. 4.**
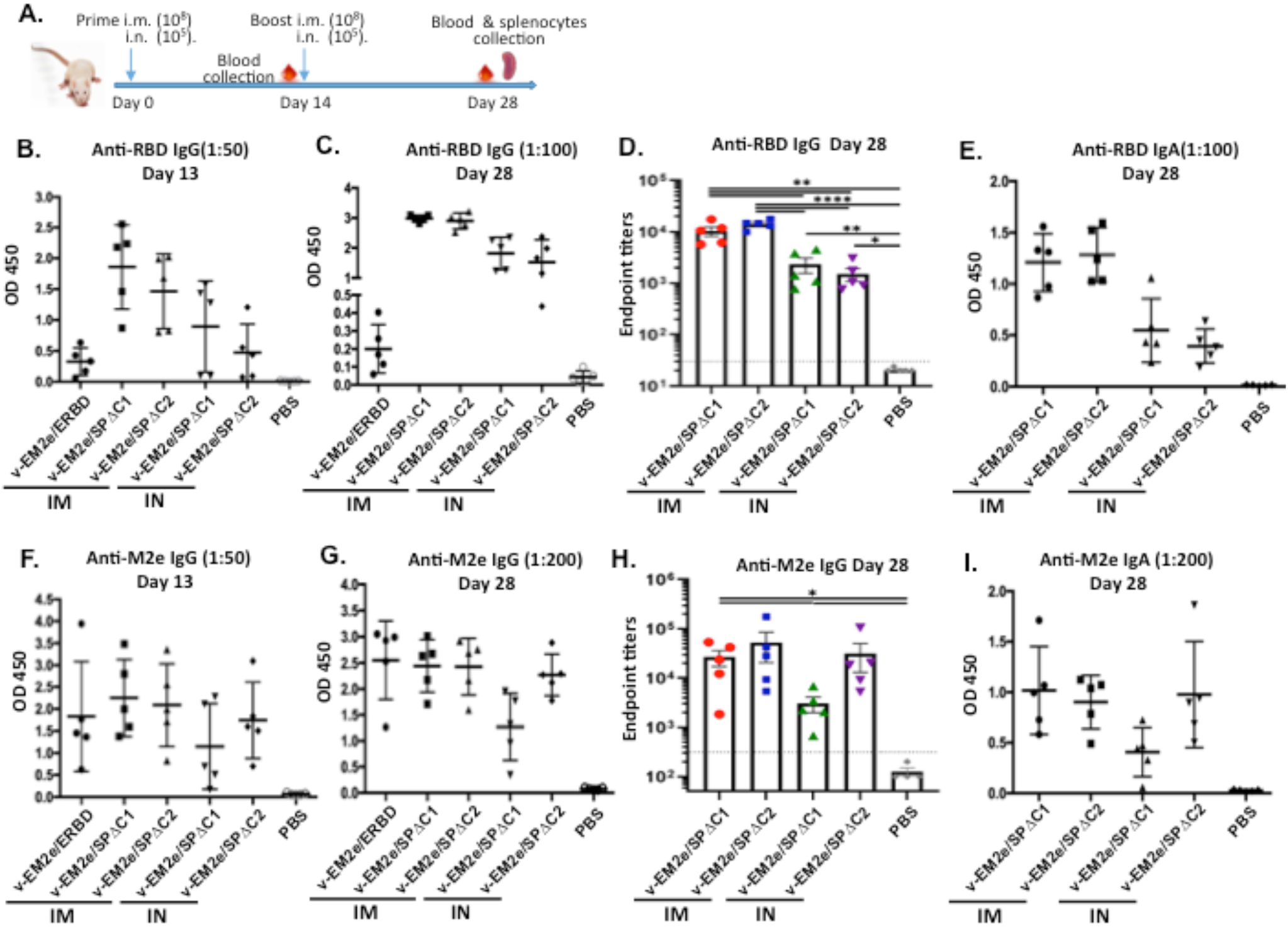
Anti-SARS-CoV-2 RBD and anti-influenza M2e immune responses induced by immunization with different bivalent VSV vaccine candidates. A) Schematic of the bivalent rVSV vaccine candidates immunization protocol in mouse. Balb/c mice were immunized with V-EM2e/SPΔC1, V-EM2e/SPΔC2 or V-EM2e/ERBD via intramuscular (IM) or intranasal (IN) routes, as indicated. The mice sera were collected at day 13 and 28 and were measured for anti-SARS-CoV-2 RBD IgG and IgA antibody levels (B-D) or measured for anti-M2e IgG and IgA antibody levels (F-H). E, D) The anti-SARS-CoV-2 RBD and anti-M2e IgA antibody levels at 28 day. Data represent Mean ±SD. Statistical significance was determined using unpaired T-test. *, P < 0.05; **, P < 0.01; ***, P < 0.001; ****, P < 0.0001.

### Vaccination with bivalent VSV vaccine candidates induced potent neutralizing antibodies that protect against infection with various SARS-CoV-2 SP pseudoviruses

The control of SARS-CoV-2 transmission relies on herd immunity among the human population, which can be obtained via infection-induced or vaccination-induced immunity. An ideal COVID-19 vaccine must be able to prevent SARS-CoV-2 infection by inducing a high level of neutralizing antibodies (nAbs). In this study, we tested whether the antibodies induced by different rVSV-based SP vaccine candidates could neutralize SARS-CoV-2 pseudoviruses (PVs) and prevent infection. Various single-round SPΔC-pseudotyped lentiviruses expressing firefly luciferase (Luc) were produced by co-transfecting 293T cells with HIV pNL4.3-/Env-Vpr-/Luc+ vector ^37^ and SPΔC-expressing plasmids, including the SPΔC of wild type (Wuhan-Hu-1), Delta (B1.617.2), Lineage B.1.617, or Beta’, which only contains the RBD mutations (K417N, E484K, N501Y and D614G) in the Beta variant. Each produced SPΔC-PV was titrated, and the appropriate amounts of PVs were used to perform the neutralizing assay in the presence of 2× serially diluted sera pooled from the mice immunized with the same vaccine candidate through the same route (2 weeks post-booster), as described in the Materials and Methods. From the results presented in Fig. 5, we found that 1) among the three IM immunization groups, the sera from mice immunized with V-EM2e/SPΔC1 contained the highest levels of neutralizing antibodies against SpΔC_WT_- or SpΔC_Delta_-PVs infections; however the V-EM2e/ERBD immunization only induced significantly lower neutralizing activity, which was consistent with the low level of anti-RBD IgG (Fig. 5A and B) and indicated that the full-length SP was more efficient in stimulating neutralizing antibody production. 2) The results also suggested that V-EM2e/SPΔC1- and V-EM2e/SPΔC2-immunized mouse sera had higher neutralizing activity against SpΔC_Delta_-PVs than against SpΔC_WT_-PVs (Fig. 5, compare B to A). Additionally, regardless of the route of administration (IM or IN), V-EM2e/SPΔC1 vaccination induced antibodies that were able to neutralize both SpΔC_B.1.617_- and SpΔC_Beta’_-PV infections (Fig. 5D and E). 3) We also noticed that the neutralizing antibody titers in sera from the IM-vaccinated groups were significantly higher than those in the IN-vaccinated groups. These results were expected because the immunization dose for the IM groups was 1000-fold higher than that for the IN groups. 4) All tested immune sera did not show any neutralization activity against VSV-G-PV infection (Fig. 5C). In summary, V-EM2e/SPΔC1, which contained full-length Delta-SPΔC, elicited high titers of neutralizing antibodies specifically against Delta SpΔC-PVs and other SARS-CoV-2 SP-PVs.

**Figure 5.**
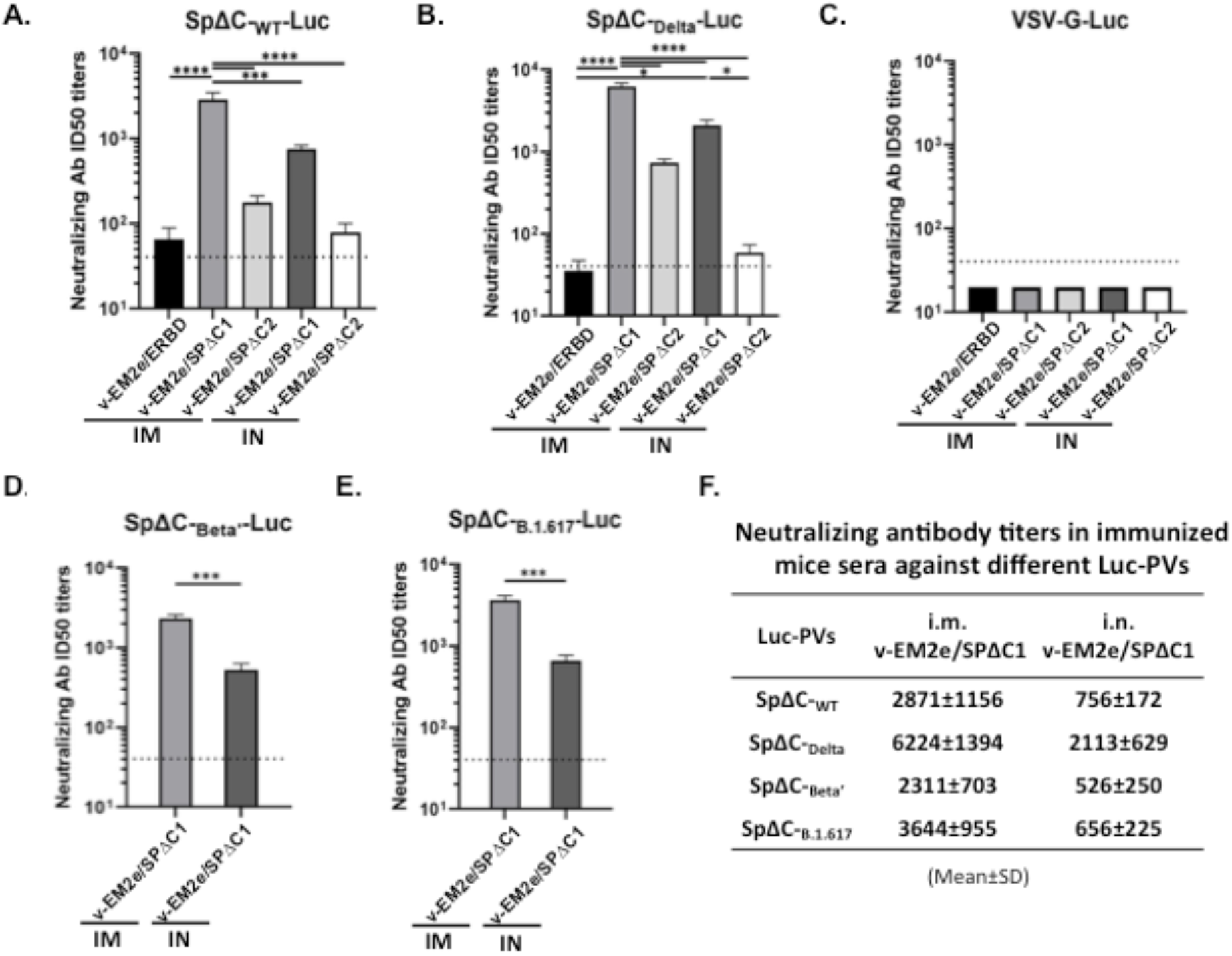
rVSV Delta SP vaccine candidates elicited neutralization antibodies. The neutralization titers (50% inhibition) in immunized mice against SPΔC_wt_-Luc-PVs (A), SPΔCdelta-Luc-PVs (B), SPΔC_beta’_-Luc-PVs (D) and SPΔC_b.1.617_-Luc-PVs (E) after 28 days of prime-boost vaccination with different bivalent vaccines. VSV-G-Luc-PVPs (C) was used as negative control. The mouse serum of each immunization group collected at day 28 were pooled together, 2× serially diluted and incubated with different Luc-PVs (∼10^4^ RLU). Then, the mixtures were added in A549_ACE2_. cell cultures and the infection of PVs was determined by Luciferase assay at 48∼66 hrs post infection. The percentage of infection was calculated compared with no serum control and neutralizing titers were calculated by using sigmoid 4PL interpolation with GraphPad Prism 9.0, as described in Materials and Methods. The neutralizing titers of V-EM2/SPΔC1 immunized mouse sera against different SPΔC-Luc-PVs were listed (F). Data represent Mean ±SD and were obtained from over three independent experiments. Statistical significance was determined using ordinary one-way ANOVA test and Turkey’s test. *, P < 0.05; **, P < 0.01; ***, P < 0.001; ****, P < 0.0001.

### Induction of Th cytokines in splenocytes from the mice immunized by the bivalent VSV vaccine candidates

Effective vaccination involves induction of T helper cells that produce cytokines to shape subsequent humoral adaptive immune responses. We therefore collected splenocytes from the naïve animals and the immunized animals (via IM or IN injection route). They were cultured in the absence of any peptides (Fig. 6A-E), with the SARS-CoV-2 SP subunit 1 (S1) overlapping peptide pool (Fig. 6F-J) or with influenza M2e peptides (Fig. 6K-O). We examined the levels of specific cytokine production in the splenocyte culture medium to determine whether T cells were stimulated in the immunized mice.

**Figure 6.**
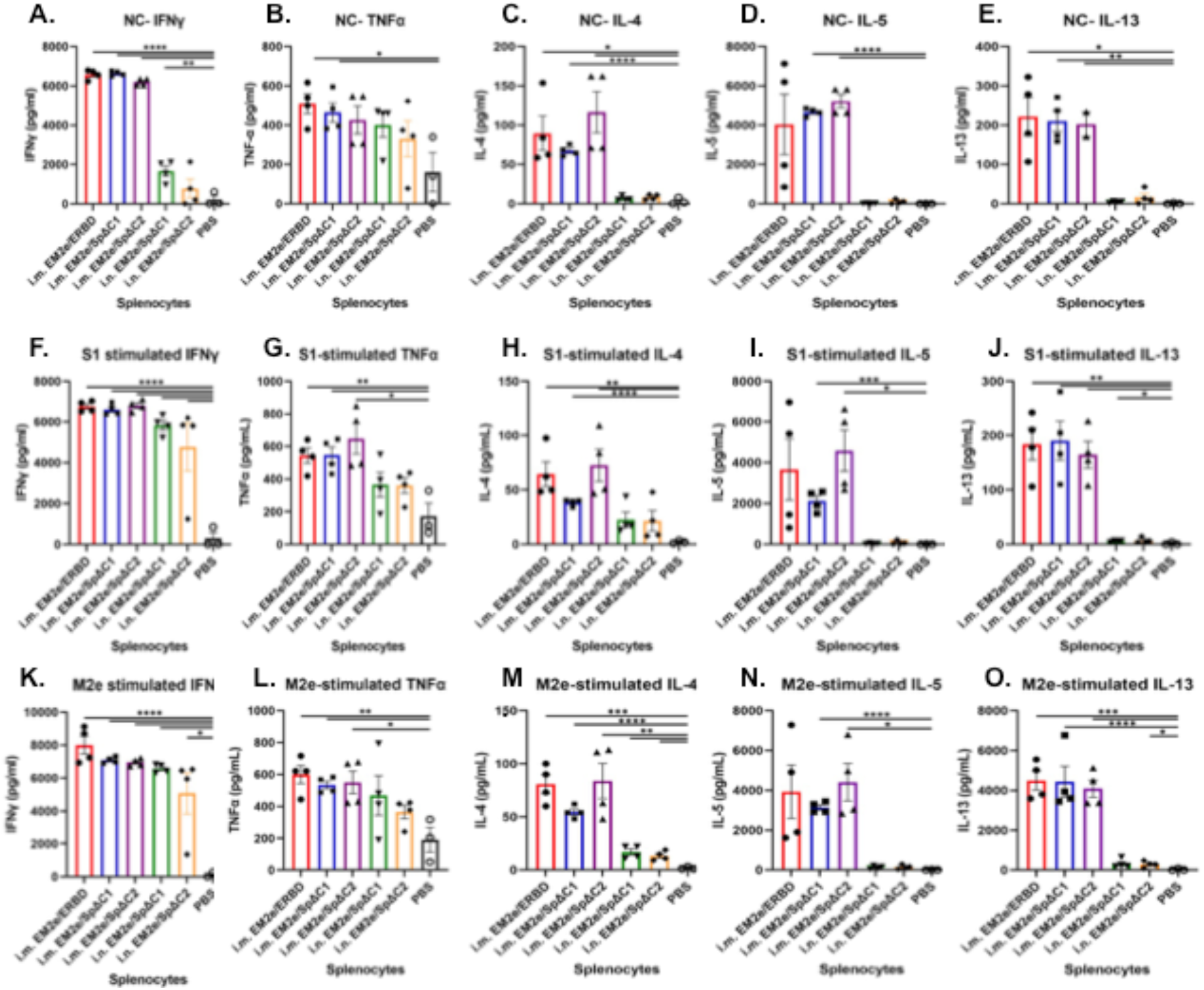
T-cell immune response induced by bivalent VSV vaccine candidates. Splenocytes isolated from immunized mice were stimulated with or without SARS-CoV-2 SP subunit 1 peptide pool or influenza M2e peptide. After 4 days of stimulation, supernatants were collected, and the release of Th1 (IFN-ϒ, TNF-α) and Th2 (IL-4, IL-5, IL-13) cytokines in the supernatants was quantified with an MSD U-plex mouse cytokine immunoassay kit and counted in the MESO Quickplex SQ120 instrument. Statistical significance between the two groups was determined using an unpaired t test. *, P < 0.05; **, P < 0.01.

As expected, we observed low/no level of Th1 cytokines (IFNγ and TNFα) and Th2 cytokines (IL-4, IL-5 and IL-13) in the splenocytes of PBS-treated control animals. In contrast, high level of Th cytokines were detected in the animals that were immunized with our vaccine candidates. For the IN-immunized mice splenocytes, slightly elevated Th1 cytokines (IFNγ and TNFα) were detected without peptide treatment, while the Th2 cytokines (IL-4, IL-5 and IL-13) in these groups were as low as those of the PBS group (Fig. 6A-E, compare bars 4-5 to bar 6). However, the stimulation with S1 or M2e peptides remarkably elevated the secretion of IFNγ and, to a lesser extent, IL-4 from IN-immunized splenocytes compared with PBS control (Fig. 6, compare F and H, to A; K and M to C, bars 4 and 5), suggesting the S1/M2e-specific re-activation ability (memory) of these splenocytes. Further, the ratios of IFNγ (Th1) and IL-5 (Th2) of these IN-immunized mice after peptide-treatments were 25∼750 (Fig, S2D and G, bars 4-5), implying that IN immunization stimulated a very strong Th1-biased response.

For IM-immunized mice, both Th1 cytokines (IFNγ and TNFα) and Th2 cytokines (IL-4, IL-5 and IL-13) showed a statistically significant elevation in the splenocytes supernatants compared with those of the naïve PBS control group. Interestingly, such high levels of cytokine production were observed in the splenocytes cultures that had no peptide re-stimulation (Fig. 6A-E, compare bars 1-3 to bar 6). Addition of specific Ag (S1, M2e peptides) could not further augment the cytokine production in the splenocytes in vitro (Fig. 6A-E, bars 1-3 to F-O, bars 1-3). The ratios of Th1 cytokines and Th2 cytokines (such as IFNγ/IL-5 and TNFα/IL-4) of these IM-immunized mice were around 1∼15 (Supplement Fig. S2A-I, bars 1-3), indicating that the IM immunization stimulated a more Th1/Th2-balanced cellular response with a little Th1-bias.

Collectively, above results suggested that our VSV-based vaccine candidates have good immunogenicity to elicit strong T-cell immune responses, and the administration via IM route can induce a Th1-biased but relatively Th1/Th2-balanced immune response while the administration via IN route will induce a very strong Th1-biased response.

### Immunization with V-EM2/SPΔC1 protects mice from lethal H1N1 and H3N2 influenza virus challenge

The above studies have demonstrated the strong humoral and cellular immune responses induced by V-EM2e/SPΔC1. We next investigated whether V-EM2e/SPΔC1 immunization could protect against influenza virus infection. Briefly, groups of 5 mice were vaccinated with V-EM2/SPΔC1 via either IM- or IN route and boosted on Day 14, while control mice received only PBS (IN) (Fig. 7A). For H3N2 challenge experiment, all mice were immunized with V-EM2/SPΔC1 via IN route, with (DD) or without (SD) boost on Day14. On Day 28, we confirmed before challenge that high levels of anti-M2e and anti-RBD antibodies were induced in all immunized mice in both vaccine delivery routes (Fig. 7B and C). Then, all mice were challenged with a fatal dose of the A/Puerto Rico/8/34 H1N1 strain (2.1×10^3^ PFU/mouse) or H3N2 (1.4×104 PFU/mouse) intranasally as previously described ^21^. Following challenge with either H1N1 strain or H3N2, a high morbidity rate was observed among the PBS group mice, exhibiting significant weight loss until death or reaching the end point (over 20% weight loss) within 5 or 6 days. (Fig. 7D and H). In contrast, both the IM and IN groups of V-EM2/SPΔC1-vaccinated mice showed slight weight loss (∼12%) until Day 4 and then almost fully recovered at Days 8-10. The survival curve further indicated that both IM- and IN-immunized mice had 100% protection against H1N1 and H3N2 infections (Fig. 7E and H). We also monitored the H1N1 or H3N2 viral loads in the lungs of infected mice. The results showed that the titer of H1N1 or H3N2 virus in PBS-treated mice reached approximately 3×10^6^ TCID50/gram or 5×10^8^ TCID50/gram of lung tissue 5 days post-challenge, while the virus titers in the immunized mice were only approximately 3×10^3^ TCID50/gram or 5×10^4^ TCID50/gram of lung tissue respectively, indicating that H1N1 and H3N2 virus replications were considerably suppressed in the lungs of immunized mice (Fig. 7F and I). Interestingly, the results also indicate that a single-dose administration via IN achieved similar protection efficiency as that double-dose administration. Overall, these results provide strong evidence that both IM and IN vaccination with the bivalent vaccine V-EM2e/SPΔC1 were safe and effectively protected mice from lethal H1N1 and H3N2 influenza challenge.

**Figure 7.**
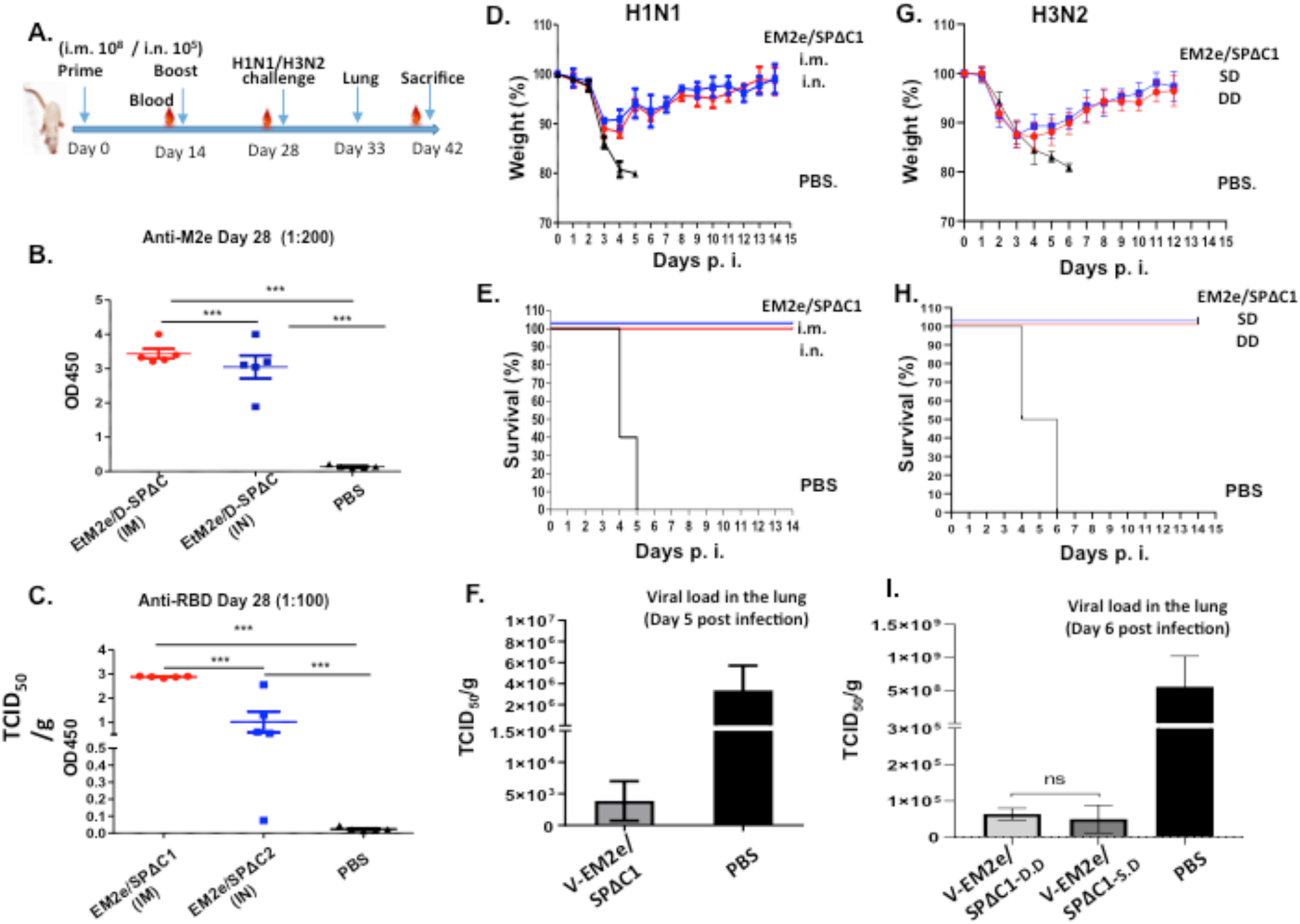
Mice immunized with V-EM2/SPΔC1 were protected against the lethal challenge of H1N1 influenza virus. A) Schematic of the bivalent VSV vaccine candidate immunization and influenza virus challenge protocol used in the study. For H1N1 challenge experiment, the Balb/c mice were immunized with 1×10^8^ TCID_50_ (IM) or 1×10^5^ TCID_50_ (IN) of V-EM2e/SPΔC1 or PBS at day 0 and day 14. On day 27, the blood samples were collected and measured for anti-SARS-CoV-2 RBD and anti-influenza M2e antibody levels by ELISA (B, C). At day 28, all the mice were challenged with 2100 PFU of H1N1 influenza virus. Weight loss (D) and survive rates (E) of the mice were monitored daily for 2 weeks. F) Viral loads in the lung tissue of immunized mice and PBS group at day 5 post H1N1 challenge were measured in MDCK cell line, as described in Materials and Methods. For H3N2 challenge experiment, the Balb/c mice were immunized with 1×10^5^ TCID_50_ (IN) of V-EM2e/SPΔC1 or PBS at day 0 (single-dose, SD), and day 14 (double-dose, DD). At day 28, all the mice were challenged with 1.4×10^4^ PFU of H3N2. G) Weight loss; H) Survive rates; I)Viral loads in the lung tissue of immunized mice and PBS group at day 6 after H3N2 challenge.

## Discussions

To date, the Delta variant is the most transmissible SARS-CoV-2 virus, and the current vaccines only provide partial protection against Delta variant infection. To rapidly respond to this critical situation, we have developed vaccine candidates that specifically target the SARS-CoV-2 Delta variant and influenza viruses (Fig. 1). These bivalent rVSV vaccines simultaneously express an EboGPΔM-M2e fusion protein (EM2e) and an EboGPΔM-SARS-CoV2 RBD fusion protein (ERBD), a full-length SPΔC_Delta_ (SPΔC1), or an S2-deleted SPΔC_Delta_ (SPΔC2). Replication kinetics studies revealed that the replication of these rVSV viruses was significantly attenuated compared to wild-type VSV and unable to infect CD4+ T lymphocytes, suggesting highly attenuated characteristics of these rVSV-based vaccines. Interestingly, our immunization study and neutralizing analysis revealed that the sera from V-EM2e/SPΔC1- and V-EM2e/SPΔC2-immunized mice exhibited significantly higher neutralizing activity against SARS-CoV-2 SP_Delta_-PV infection than against SP_WT_-PV infection (Fig. 5). Unfortunately, rVSV-EM2e/ERBD immunization did not elicit either high levels of antibody responses or neutralizing activity. To date, the reasons for the low antigenicity of RBD fused with EboGPΔM in our system remain unknown, since EboGPΔM-M2e still induced robust immune responses (Fig. 4). Interestingly, several previous studies have demonstrated that RBD vaccine candidates were able to induce sufficient neutralizing antibody (nAb) responses and provided some protection against SARS-CoV-2 ^38^. The Inconsistencies in immunogenicity of RBD in different studies may be due to the difference in the delivery method of RBD, such as in nanoparticles or in rVSV, and deserves further study.

An important unique feature of the vaccine platform in this study is that it can simultaneously protect against SARS-CoV-2 and influenza viruses. In particular, this vaccine is designed to specifically target the SARS-CoV-2 Delta strain and the highly conserved ectodomain of M2 (M2e) of influenza virus that were derived from human, avian and swine virus strains ^39^ (Fig. 1A, d). Our recent study demonstrated that rVSV expressing EboGPΔM-M2e alone induced robust anti-M2 humoral responses and effectively protected mice against either H1N1 or H3N2 virus challenge ^21^, indicating the great potential of rVSV-EboGPΔM-M2e as a universal influenza vaccine. In this study, the results revealed that all vaccine candidates elicited a high level of M2-specific immune response. We further demonstrated that the lead vaccine candidate V-EM2/SPΔC1, via either IM or IN route, effectively protected mice from lethal H1N1 influenza virus infection. As for protection against SARS-CoV-2, the serum samples from vaccinated mice displayed robust antibody responses and high neutralizing activity against SARS-CoV-2 SP-PV infections. Of course, more studies, especially animal experiments, are required to demonstrate protection against SARS-CoV-2, especially the Delta variant, and other strains of influenza. Nevertheless, the data presented here suggest that the V-EboGP-ΔM-based multivalent vaccines may represent an attractive approach to respond to the COVID-19 and influenza pandemics. Since V-EboGP-ΔM can be used as a vector to express heterologous polypeptide antigens and/or epitopes of pathogens, it will also allow us to develop vaccines against other pathogens during future epidemic/pandemic outbreaks.

The attenuated replication-competent VSV vaccine is an ideal platform for developing novel vaccine candidates against outbreak pathogens ^40^. In addition to its safety and easy and scalable production, the VSV vaccine induces a rapid and robust immune response to viral antigens after a single immunization and has been shown to protect against several pathogens ^23-27^. Currently, several VSV-based SARS-CoV-2 vaccines have been reported ^41-45^. By expressing the SARS-CoV-2 spike gene in place of the VSV glycoprotein gene, several studies have shown that a single-dose vaccination resulted in a rapid and potent induction of SARS-CoV-2 neutralizing antibodies and protected mice or hamsters against SARS-CoV-2 challenge. Interestingly, immunizing rhesus macaques with the VSV vaccine expressing the SARS-CoV-2 spike protein and the EBOV glycoprotein (VSV-SARS2-SP-EBOV) via IM but not IN offered rapid protection against COVID-19 ^45^. In agreement with the above observation, our study also showed that IM administration of V-EM2e/SPΔC1 or V-EM2e/SPΔC2 induced higher titers of neutralizing antibodies and more robust cellular immune responses than IN administration. However, it should be noted that the dose of VSV vaccine used for IN administration was 1000-fold lower than that for IM administration, suggesting the possibility to achieve stronger immune responses if a higher dose of VSV vaccine is administered via the IN route.

The rapid T cell response following vaccination is a key part of the immune response required to elicit effective protection. In this study, we observed strong T cell responses such as high levels of secreted cytokines, including IFN-γ and IL-4, in splenocytes from mice stimulated with rVSV-based vaccine candidates. These findings implied that rVSV-based vaccines could elicit both humoral and cellular responses in mice. Interestingly, we found a significant difference in the ratios of Th1-cytokine/Th2-cytokine (IFN-γ/IL-5) between the samples isolated from the IM- and IN-immunized mice, suggesting that IM immunization triggered a Th1-biased but relatively Th1/Th2-balanced cellular response, while IN immunization induced an extremely Th1-biased cellular response (Supplement Fig. S2). Th1 cells are responsible for the activation of B cells (producing IgG2a), macrophages, and cytotoxic T cells, and Th2 cells mostly activate B cells (producing IgG1 and IgE). Furthermore, secreted Th1 cytokines, such as IFN-γ and TNF-α, tend to induce pro-inflammatory reactions, whereas secreted Th2 cytokines, such as IL-4, IL-5 and IL-13, play anti-inflammatory roles to suppress excessive inflammation. Therefore, a balanced Th1/Th2 response is critical for the host to clear the virus and avoid damages caused by uncontrolled inflammation. Given that Th2 responses may be associated with antibody-dependent enhancement (ADE) of infection in mice vaccinated with inactivated vaccines and challenged by SARS-CoV-1 ^46^, it is encouraging that a Th1-biased cellular response was elicited by our rVSV-based SARS-CoV-2 vaccines regardless of the administration route. However, the effect of different delivery routes on the protective efficacy of the vaccine is currently unclear. Compared with more invasive IM immunization, the IN immunization process more closely resembles natural infection with SARS-CoV-2. Hence, more detailed studies are required to determine whether IN administration will enable the vaccine to establish more effective protection.

The safety profile is also an important issue for vaccine development. Even though the pathogenicity of the rVSVΔG vector is significantly attenuated compared to the wild-type VSV, the replacement of EboGP-ΔM and SARS-CoV-2 SP affected the cell tropism of vaccine candidates. As expected, we observed much attenuated replication kinetics of rVSV-based vaccines in various cell lines compared to rVSV expressing VSV-G. Except for Vero-E6 cells, these vaccines showed no or much milder cytopathic effects in most tested cell lines compared with VSV_wt_, which induced significant cytopathic effects (Fig. 3). Importantly, these rVSV-based vaccines do not target CD4^+^ T cells, which is also essential for protecting the immune system from attack. Given the fact that SARS-CoV-2 SP_Delta_ was able to efficiently mediate cell fusion and subsequently caused the cell death ^32 47^, we introduced a mutation (I742A) into the SPΔC_Delta_ gene of V-EM2e/SPΔC1 to reduce SP_Delta_’s cytotoxicity. Indeed, the TI742A mutation significantly reduced SPΔCDelta-pseudovirus’s infectivity and syncytia formation (Fig. 1B-D). Also, in V-EM2e/SPΔC2, we deleted a 381 aa sequence encompassing the S2 domain of the SP_Delta_ (Fig. 1A,b). In comparison of immunogenicity of these two vaccine candidates, both V-EM2e/SPΔC1 and V-EM2e/SPΔC2 elicited similarly high levels of anti-SARA-CoV-2 IgG and IgA antibody responses (Fig. 4) and induced comparable levels of T cell cytokine production (Fig. 6). However, the level of neutralization antibody induced by V-EM2e/SPΔC2 was not as high as that of V-EM2e/SPΔC1. (Fig. 5A and B), This may be because the S2 domain deletion removed some fusion-related regions that are important for inducing neutralizing antibodies. Indeed, a previous report indicated that two segments, 884-891 and 1116-1123, located in SP-S2, are very effective in inducing host immune responses ^48^. It should be noted that a recent report clearly indicated that a VSV-vectored vaccine expressing SARS-CoV-2 S1 alone protected against severe COVID-19 virus infection in a hamster model ^42^. Therefore, whether V-EM2/SPΔC2 could achieve effective protection against SARS-CoV-2 Delta variant infection is still an open question waiting for investigation.

Overall, in this study, we generated rVSV-based bivalent vaccines and demonstrated that immunization with these rVSV-vectored vaccines in mice induced sufficient immune responses, including neutralizing antibodies and/or cell-mediated immune responses against SARS-CoV2 Delta variants and influenza virus. Our study also showed that immunization with the lead vaccine candidate V-EM2e/SPΔC1 via both IM and IN effectively protected mice from lethal H1N1 and H3N2 influenza virus infection. Further investigation of its efficacy to protect against SARS-CoV-2 Delta variants and more influenza strains in animal models, including in non-human primates, will provide substantial evidence for new avenues for controlling two contagious respiratory infections, COVID-19 and influenza.

## Materials and methods

### Plasmid constructions

In this study, the gene encoding SPΔC_Delta_ was amplified from the previously described plasmid pCAGGS-SPΔC_Delta_ ^32,^ and the I742A mutation was introduced by site-directed mutagenesis technique with, 5’-primers 5_TGTACAATGTATGCATGCGGAGACAGC, and 3’-primer, 5_GCTGTCTCCGCATGCATACATTGTACA. Then, the amplified SPΔC_Delta_-I742A gene was cloned at *XhoI* and *NheI* sites of an rVSV-based influenza vaccine vector, rVSV-EΔM-M2e ^21^, and the constructed plasmid was named rVSV-EM2e/SPΔC1. To construct rVSV-EM2e/SPΔC2, we used a two-step PCR technique to generate cDNA that carried an additional 381 aa deletion in the S2 region of SPΔC_Delta_ (Fig. 1A, b), and the amplified SPΔC2-encoding cDNA was also cloned into the rVSV-EΔM-M2e vector using the same restriction enzymes, yielding rVSV-EM2e/SPΔC2. To construct rVSV-EM2e/ERBD, a cDNA fragment encoding the receptor binding domain (RBD) of SARS-CoV-2 (Wuhan-Hu-1, GenBank accession No. MN908947) spike protein was amplified from a pCAGGS-nCoVSP plasmid ^31^ and inserted in pCAGGS-EboGPΔM at the MLD region ^21^. Then, the EboGPΔM-RBD cDNA (Fig. 1A, d) was cloned into the *XhoI* and *NheI* sites of the rVSV-EΔM-EM2e vector and named rVSV-EM2e/ERBD (Fig. 1B). To construct pCAGGS-SPΔCBeta’ and pCAGGS-SPΔC_B.1.617_, the mutations K417N, E484K, N501Y and D614G (for SPΔCBeta’) and L452R, E484Q and D614G (for SPΔC_B.1.617_) were introduced into the pCAGGS-nCoVSPΔC plasmid ^32^ by site-directed mutagenesis. All rVSV vectors and inserted transgenes were confirmed by sequencing.

### Cells, antibodies, chemicals and viruses

A human embryonic kidney cell line (HEK293T), a human lung type II pulmonary epithelial cell line (A549), a human fetal lung fibroblast cell line (MRC-5), a human glioblastoma-derived cell line (U251GM), VeroE6 and MDCK cell line were cultured in Dulbecco’s modified Eagle’s medium, minimum essential medium (MEM) or DMEM/F-12 medium (21331-020, Gibco). CD4^+^ Jurkat cells were cultured in RPMI-1640 medium. A549 cells expressing the ACE2 receptor (A549_ACE2_) were described previously ^32^. Human monocyte-derived macrophage MDMs and dendritic cells (MDDCs) were prepared from human peripheral blood mononuclear cells (hPBMCs) isolated from healthy donors following procedures as described previously ^49^. All cell lines were grown in cell culture medium supplemented with 10% fetal bovine serum (FBS), 1× L-glutamine and 1% penicillin and streptomycin. The antibodies used in the study included the rabbit polyclonal antibody against SARS-CoV-2 SP/RBD (Cat# 40150-R007, Sino Biological), anti-SARS-CoV-2 S-NTD antibody (E-AB-V1030, Elabscience), anti-M2 monoclonal antibody (14C2: sc-32238, Santa Cruz Biotech.), and anti-VSV-Nucleoprotein, clone 10G4 (Cat# MBAF2348, EMD Millipore Corp). The HIV-1 p24 ELISA Kit was obtained from the AIDS Vaccine Program of the Frederick Cancer Research and Development Center. Recombinant SARS-CoV-2 proteins or peptides used in this paper include S1-RBD peptides (RayBiotech, Cat# 230-30162) and S1 overlapping peptide pool (JPT Peptide Technologies, Germany, Cat# PM-WCPV-S-SU1-1; 166 peptides; 15mers with 11 aa overlap). Influenza M2e peptide and mouse-adapted strain A/Puerto Rico/8/34 (H1N1) were described previously ^21^.

### VSV rescue and virus growth kinetics experiments

Replication-competent rVSV was recovered in 293T-Vero E6 co-cultured cells as described previously ^21^. All three bivalent VSV vaccine candidates were propagated and titrated on Vero E6 cells. To examine the growth kinetics of bivalent rVSV, cell lines were grown to confluency in a 24-well plate and infected in duplicate with VSVwt, V-EM2e/SPΔC1, rV-EM2e/SPΔC2 or V-EM2e/ERBD at a dose of 100 TCID50. After 2 hrs of incubation, the cells were washed and cultured in DMEM or RPMI containing 2% FBS. The supernatants were collected at 24, 48, 72 and 96 hours. The titers of rVSV in the supernatant were determined by the TCID50 method on Vero E6 cells in 96-well plates.

To detect the expression of EM2, Delta SPΔC, RBD, and other viral proteins in cells, rVSV-infected cells were lysed and analyzed by SDS–PAGE and WB with anti-M2e (14C2), anti-SARS-CoV-2-RBD, or anti-VSV N antibodies.

### Immunofluorescence assay and syncytia formation assay

As previously described ^21^, Vero E6 cells were grown on glass coverslips (12 mm^2^) in 24-well plates and infected with V-EM2e/SPΔC1, V-EM2e/SPΔC2 or V-EM2e/ERBD for 48 hours. After infection, cells on the coverslip were fixed with 4% paraformaldehyde for 15 minutes and permeabilized with 0.2% Triton X-100 in PBS. The glass coverslips were then incubated with primary antibodies specific for M2e or SP/RBD followed by corresponding FITC-conjugated secondary antibodies. Cells were viewed under a computerized Axiovert 200 inverted fluorescence microscope (Zeiss).

To test SARS-CoV-2 SP_Delta_-mediated syncytia formation, 293T cells were transfected with various SPΔC plasmids using Lipofectamine 2000. After 24 hrs, the cells were washed, resuspended and mixed with A549_ACE2_ cells at a 1:3 ratio and plated into 12-well plates. At different time points, syncytium formation was observed, counted and imaged by bright-field microscopy with an Axiovert 200 fluorescence microscope.

### Virus production and infection experiments

SARS-CoV-2 SPΔC-PVs (SPΔCwt-, SPΔC_Delta_-, SPΔC_Delta_-a742-PVs) were produced by co-transfecting 293T cells with each of the pCAGGS-SPΔC plasmids, pCMVΔ8.2 and Gluc expressing HIV vector ΔRI/E/Gluc ^31^. After 48 hrs of transfection, cell culture supernatants were collected, VPs were purified and quantified by HIV-1 p24 amounts using an HIV-1 p24 ELISA, as described previously ^32^. To measure the infection of SPΔC-pseudotyped VPs, equal amounts of each SPΔC-PV (as adjusted by p24 levels) were used to infect A549_ACE2_, the supernatants were collected, and the viral infection levels were monitored by measuring Gaussia luciferase (Gluc) activity.

### Cell-based pseudovirus neutralization assays

Different SARS-CoV-2 SP pseudoviruss expressing luciferase were prepared and titrated as follows: HIV-based SARS-CoV-2 SP pseudoviruses (PVs) expressing luciferase (Luc) were produced by co-transfection of 293T cells with an HIV vector (pNL4-3-R-E-Luc) ^37^ and each pCAGGS-SPΔC or pCAGGS-VSV-G plasmid by using polyetherimide (PEI) transfection in a 6-well plate. The supernatants were harvested at 72 h post-transfection, passed through a 0.45 μm filter, aliquoted and titrated on A549 cells expressing human ACE2 (A549/hACE2). The pseudovirus neutralization assay was performed on A549/hACE2 cells according to previously reported methods with some modifications ^50, 51^. Briefly, inactivated mouse sera of the same experimental group were pooled together. SPΔC pseudotyped Luc-PVs (PV-Luc-SpΔC) and control VSV-G-pseudotyped Luc-PV-Luc (25 *μ*L, ∼104 RLU) were pre-incubated with 2× serially diluted mouse sera (25 *μ*L) in a 96-well plate for 1.5 h at room temperature with gentle shaking. Then, A549/hACE2 (1.25×10^4^ cells/well, 50 *μ*L) and polybrene (final conc. 5 *μ*g/mL) were plated in the above wells containing a mix of pseudovirus and sera. After gentle shaking for 10 min, the cells were incubated at 37 °C overnight. The supernatant was removed the next day, and fresh medium was added. At 48∼66 hrs post-infection, cells were washed and lysed, and the luciferase RLU in lysate was measured. The RLU percentages (%) were obtained by comparing the RLU of tested samples to the RLU of the control samples after subtraction of the background RLUs (cell only/virus only). The neutralizing titers or half-maximal inhibitory dilution (ID50) were defined as the reciprocal of the serum maximum dilution that reduced RLU by 50% relative to no-serum (virus and cell) controls. The ID50 was calculated by using sigmoid 4PL interpolation with GraphPad Prism 9.0. All data were from at least three experiments and are shown as the means ± standard error of the means (SEMs).

### Mouse immunization and viral challenge

Female BALB/c mice aged 4–6 weeks used in this study were obtained from the Central Animal Care Facility, University of Manitoba (with animal study protocol approval No. 20-034). For rVSV immunization, mice (five per group) were immunized intramuscularly (IM, 1×10^8^ TCID_50_) or intranasally (IN, 1×10^5^ TCID_50_) with rVSV vaccine candidates on Day 0 and boosted on Day 14. Mice were sacrificed on Day 28, and spleens were harvested. Blood samples were collected on Days 13 and 28.

For influenza virus challenge in mice, the mouse-adapted strain A/Puerto Rico/8/34 (H1N1) was used. Three groups of mice (5 for each group) were IM-immunized with 1×10^8^ TCID_50_ or IN-immunized with 1×10^5^ TCID_50_ of V-EM2e/SPΔC1 or PBS on Day 0 and boosted on Day 14. On Day 28, all the mice were intranasally infected with H1N1 virus (2.1×10^3^ PFU/mouse) or with H3N2 virus (1.4×10^4^ PFU). Weight and survival of the mice were monitored daily for 2 weeks after the challenge. Additionally, 5 to 6 days post-challenge, the mice from the PBS group and two mice from the vaccination group were sacrificed, and the lungs were collected and immediately stored at −80 °C. The lung was homogenized using a tissue grinder and centrifuged at 5,000 rpm. The supernatant was used for titration in MDCK cells according to the method described previously ^52, 53^.

### Enzyme-linked Immunosorbent Assay (ELISA) for measurement of anti-SARS-CoV-2-SP/RBD or anti-influenza M2e antibody levels in immunized mouse sera

Anti-SARS-CoV-2-SP/RBD antibodies and anti-influenza M2 antibodies in mouse sera were determined by ELISA, as previously described with some modifications ^21^. Briefly, ELISA plates (NUNC Maxisorp, Thermo Scientific) were coated with 100 μl of recombinant RBD protein or M2e peptide (0.75 μg/ml or 0.5 μg/ml, respectively) in coupling buffer (0.05 M sodium carbonate-bicarbonate, pH 9.6) overnight at 4 °C. Then, the different dilutions of serum samples were incubated in the coated plate for 2 hrs at 37 °C followed by the addition of horseradish peroxidase-conjugated goat anti-mouse IgG or IgA for 1 h at 37 °C. Finally, TMB (Mandel Scientific) was added, and the absorbance at 450 nm (OD450) was measured ^54^. To determine the endpoint titers, 100 μl of 3× serially diluted sera was used to measure the OD450. The endpoint titer is designated as the reciprocal of the highest dilution of a serum that has an OD450 above the cutoff (10× negative control) and is calculated by using sigmoid 4PL interpolation with GraphPad Prism 9.0.

### Vaccine candidates-induced T cell responses

The mice were vaccinated according to the schedule described in Fig. 4A and sacrificed on Day 28 (2 weeks after booster). To assess general T cell reactivity, mouse splenocytes were collected as described previously ^29^ and plated in 48-well plates (2×10^6^/200 μl per well) in RPMI (no-peptide control) or incubated with a SARS-CoV-2 S1 overlapping peptide pool or with the influenza virus M2e peptide (1 μg/ml for each peptide). The PMA/ionomycin cocktail (Invitrogen, 81 pM/1.34 *μ*M) served as a positive control. To measure the extracellular cytokines released from splenocytes, the supernatants were collected after 4 days and stored at −70 °C. The Meso Scale Discovery (MSD) immunoassay was performed on a customized mouse U-plex Biomarker Group1 Assays kit (Mesoscale Discovery, USA) to determine the cytokines (IFN-γ, TNF-α, IL-4, IL-5 and IL-13) and analyzed on the MESO Quickplex SQ120 instrument following the manufacturer’s instructions.

### Statistics

Statistical analysis of antibody/cytokine levels was performed using the unpaired t test (considered significant at P≥0.05) by GraphPad Prism 5.01 software. The statistical analysis of neutralizing antibodies was performed using the one-way ANOVA multiple comparison test followed by Tukey’s test by GraphPad Prism.

## DATA AVAILABILITY

The data that support the findings of this study are available from the corresponding authors upon reasonable request. Source Data are provided with this paper.

## ACKNOWLEDGEMENTS

We thank Dr. Kevin Coombs for kindly providing the U251 cell lines and Mr. William Peter Stefura for his technique supports. T. Olukitibi is a recipient of the University Manitoba Graduate scholarship. This work was supported by Canadian 2019 Novel Coronavirus (COVID-19) Rapid Research Funding (OV5-170710) by Canadian Institute of Health Research (CIHR) and Research Manitoba, and CIHR COVID-19 Variant Supplement grant (VS1-175520). This work was also supported by the Manitoba Research Chair Award from the Research Manitoba (RM) to X-J.Y.

## AUTHOR CONTRIBUTIONS

The experiments were conceived of and designed by S.K., K.R.F., D.K., and X.J.Y. The constructions and rescue of different rVSV viruses and the viral replication characterization were performed by X.J.Y and Z.J.A. SARS-CoV-2 SP-pseudotyped virus infections and the neutralizing experiments were conducted by M.O and Z.J.A. Animal immunization studies were performed by M.O., Z.J.A., O.T.A., and M.L.Z., Cytokine production analyses were M.O., M.L.Z., and Z.J.A. Animal challenge experiments were carried out by O.T.A., M.O., B.W., R.V., and T.T. The initial draft of the paper was written by M.O., Z.J.A., and O.T.A. All other authors contributed to edit the paper into its final form. The work was managed and supervised by S.K., K.R.F., D.K., and X.J.Y.

